# A Systematic Review and Meta-analysis of the Protective Effect of FMDV VP1 on Animals

**DOI:** 10.1101/2020.07.23.217299

**Authors:** Peng Wu, Qingqing Liu, Jinke He, Xiaoyu Deng, Xinyue Yin, Yueli Wang, Yunfeng Zhang, Changfu Chen

## Abstract

The FMDV VP1 protein has different structures which could decrease or increase the immune response. We undertook a meta-analysis to evaluate the protective effect of VP1 on the FMDV. A systematic search of the PubMed, Embase, CNKI and Wan fang DATA was conducted up to April 2020. Experimental studies involving the VP1 protection effect on FMDV were included. Extracted data were analyzed using Rev-Man 5.3 software. Chi-square tests were used to analyze the heterogeneity among the documents. The fixed-effect model was used for meta-analysis to find the combined effect value and 95% confidence interval. Sensitivity analysis was performed on the differences in the combined values of model effects, and the inverted funnel chart method was used to assess the publication bias of the included literature. A total of 12 articles were included for meta-analysis. The results of showed that VP1 had a protective effect on FMDV [*MH* = -0.66, 95% *CI* = (−0.75, -0.56), *P* < 0.00001]. Sensitivity analysis showed that the results were robust. The funnel graph method showed that the published literature had a small publication bias and met the requirements of this study. It is necessary to study the epitopes of VP1 to produce new vaccines. VP1 could protect animals from FMDV attacks. It is necessary to study the VP1 protein and its epitopes and use it as a new vaccine and diagnostic product.

## Introduction

Foot-and-mouth disease (FMD) is one of the most important infectious diseases of cloven-hoofed livestock and wildlife^[1]^, which is an acute, hot, and highly contagious infectious disease caused by the foot-and-mouth disease virus (FMDV). In 2007, the British FMD epidemic caused a major blow to the export of cloven-hoofed animals and their products (http://www.defra.gov.uk). FMD is a major hindrance to international trade in animals and animal products, and the cost of eradication can be enormous^[2]^. For this reason, the prevention of FMD has become one of the most important public health goals today.

Vaccine immunization is an economical and effective means of controlling FMD. The attenuated FMD vaccine has played a huge role in controlling the FMD epidemic. However, due to the possibility that the vaccine strain might regain its virulence, its use has been suspended. At present, most of the vaccines used for FMD are inactivated. Although this is safer than the weakened live virus vaccine, cultivating viruses and rendering them inactive still involve considerable biosecurity risk^[3]^. In recent years, FMD synthetic peptide vaccines, genetically engineered subunit vaccines, epitope vaccines, and other new vaccines have also seen made great progress. Although FMDV vaccines can induce good humoral protective immunity, the immune response is not quick enough for use in emergency vaccination campaigns meant to stop the spread of FMD ^[4]^. The duration of immunity is also short.

FMDV belongs to the genus Aphthovirus of the Picornaviridae family^[5]^. The viral genome is about 8500 nucleotides in length. The capsid of the FMDV virus is composed of VP1, VP2, VP3, and VP4. VP1, VP2, and VP3 are partially exposed on the surface of virus particles and are all immunogenic, but only VP1 could independently produce neutralizing antibodies. VP1 protein is the most important antigen protein of FMDV. It is exposed to the surface of virus particles on the form of protrusions and is the main inducer of neutralizing antibodies^[6, 7]^. Therefore, the use of VP1 protein and antigenic determinants as new vaccines and diagnostic products has become a research hotspot.

The VP1 protein has a unique RGD motif on a flexible loop structure. This sequence is highly conserved among FMDV strains. It is the cell receptor site of FMDV. It mediates the adsorption of viruses and cells by binding to integrin receptors on the cell surface^[8]^. The G-H loop of VP1 is the main linear epitope that induces FMDV neutralizing antibodies^[9]^. Its spatial configuration is relatively complex, with relatively large movements. The synthetic FMDV G-H loop peptide has been shown to stimulate guinea pigs to produce high levels of neutralizing antibodies^[10]^, which means a good immune effect^[11]^.

Recent studies have shown that VP1 can cause the body to produce neutralizing antibodies, so it could serve as a vaccine to protect animals from FMDV. However, in our study, it was found that VP1 expressed in prokaryotic cells and VP1 expressed in recombinant vectors could not induce the body to produce neutralizing antibodies, nor could it be detected by liquid phase blocking ELISA. Some research has shown that FMDV VP1 can also suppress immune responses. Type I interferon is considered to be an important part of the innate immune response, especially for viral infections^[12]^. According to reports, VP1 inhibits tumor necrosis factor (TNF-α) and Sendai virus-induced type I interferon response in HEK293T cells. Related studies have found that sorcin, a protein that regulates the response of cells to viral infections, can interact with VP1 to suppress the type I interferon response and so play a role in suppressing the innate immune system^[13∼16]^. The FMDV VP1 has been identified as an interferon inhibitor because it interacts with soluble resistance-related calcium-binding protein (sorcin)^[14]^. Most new vaccines, especially nucleic acid vaccines and recombinant viral live vector vaccines, are built based on the VP1 protein. With these premises, we performed a meta-analysis of the currently available studies to comprehensively explore the effescts of VP1 on FMDV prevention.

## Material and methods

This systematic review was conducted in accordance with the Preferred Reporting Items for Systematic Reviews and Meta-analyses guidelines (PRISMA) (Supporting Information Table S1).

### Literature search strategy

This meta-analysis was performed on documents published before April 2020 using a computer and was searched by two researchers respectively. A computerized search for the VP1 protection effect on FMDV was conducted using the databases of The National Library of Medicine (Medline via PubMed), Embase, China National Knowledge Infrastructure (CNKI), Wan Fang DATA, using the following keywords: “FMDV,” “VP1,” “Vaccine,” and “protection,”.

### Inclusion and exclusion criteria7080

Incorporate literature standards. ➀ Published documents included Chinese-language and English-language works on FMDV VP1 protein immunized animals. ➁ VP1 could be expressed by various vectors. ➂The work must have used FMDV to carry out the attack protection experiment. Excluding literature standards. ➀ The literature refers to the research progress, review or irrelevant protection effect. ➁ The research object was not FMDV. ➂ The immunized antigen was not VP1. Literature research results did not provide the necessary basic data or provided incomplete data. ➃ Replace unusable documents such as repeated reports and poor quality ones.

### Data extraction tired

Two researchers conducted preliminary screening by reading study titles and abstracts, then read the full text and selected works for this analysis according to the inclusion and exclusion criteria. In case of different opinions, matters were resolved by discussion. We independently extracted the data and entered them into a specially designed data extraction table. Data extracted included the first author, time of publication, number of animals, number of protections, and similar information.

### Statistical analysis

The database was established using Microsoft Office Home and Student 2019 software and the review management software (RevMan 5.3) provided by the Cochrane Collaboration Network which was used for meta-analysis. Chi-square tests were used to analyze the heterogeneity among the documents. When there was no statistical heterogeneity between the documents, the fixed-effect model was used for meta-analysis to find the combined effect value and 95% confidence interval. Sensitivity analysis was performed on the differences in the combined values of model effects, and the inverted funnel chart method was used to assess the publication bias of the included literature.

## Results

### Identification of suitable studies

The process of document retrieval and screening is shown in Fig 1. A total of 1367 articles in the immune effect of VP1 were retrieved. After removing 48 duplicate articles and consulting the titles and abstracts of those that remained, a total of 59 articles were found to meet the inclusion criteria. In the included literature, 12 articles reported the immune effect.

**Fig.1.**
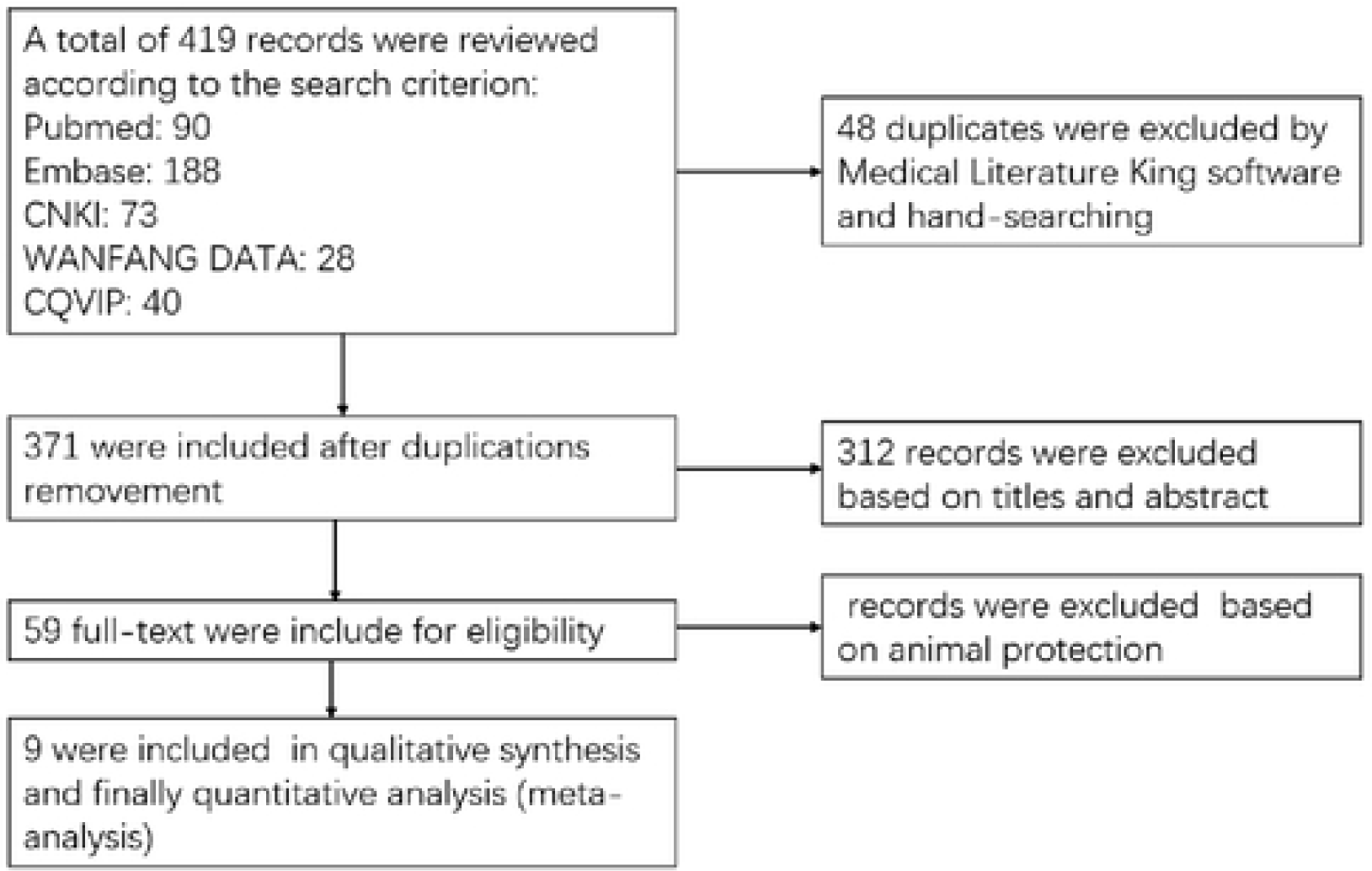
Flowchart of included and excluded trials.

### Characteristics of the reports

The characteristics of the included studies are given in Table 1. Of the included studies, nearly all were conducted in Eastern countries. The total number of included animals was 195. The total number of experimental cases included was 66. We also observed that all of the studies were conducted between 2004 and 2015.

**Table 1.** Characteristics of eligible trials.

| Study ID | Experimental |  | Control |  |
| --- | --- | --- | --- | --- |
|  | Events | Total | Events | Total |
| Huali2007 | 6 | 9 | 5 | 5 |
| Jingyun2004 | 2 | 4 | 4 | 4 |
| Li2005 | 4 | 10 | 10 | 10 |
| Li2006 | 4 | 10 | 10 | 10 |
| Manlin2008 | 3 | 26 | 18 | 18 |
| Miao2015 | 1 | 10 | 4 | 5 |
| Xiaolong2005 | 13 | 29 | 10 | 10 |
| Yong2005 | 31 | 90 | 10 | 10 |
| Zhuang 2010 | 2 | 7 | 4 | 4 |

### Meta-analysis

To correct the problem of unsatisfactory test performance caused by the small number of documents, we here used a combination of statistical values and q test. Comparing the immune effect of the control group and the VP1 experimental group, the heterogeneity analysis *I*^*2*^ =43%, used fixed effect model analysis. It is statistically significant or *I* ^*2*^< 50% think that there was no heterogeneity in the study. The analysis results showed that VP1 had a protective effect on FMDV [MH = -0.66, 95% CI = (−0.75, -0.56), *P* <0.00001](Fig 2).

**Fig 2.**
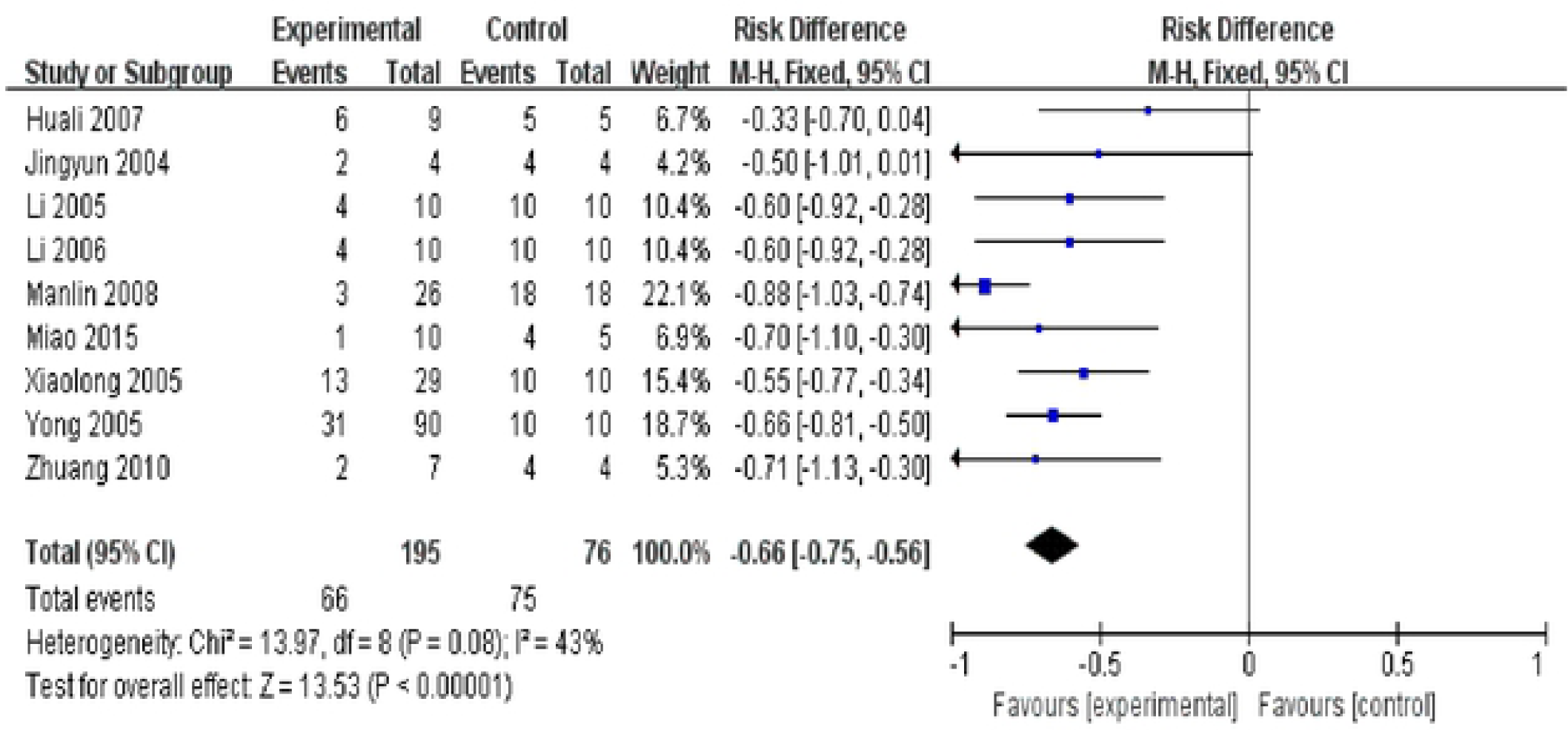
Forest plot of the protective effect of VP1 on FMDV.

### Publication bias

The funnel graph method was used to control the publication bias of the meta-analysis of selected literature. By observing the funnel graph of meta-analysis, we found that although the funnel graph was not completely symmetrical, it was still within an acceptable range (Fig 3). This indicated that the published literature has a small publication bias, which met the requirements of this study.

**Fig 3.**
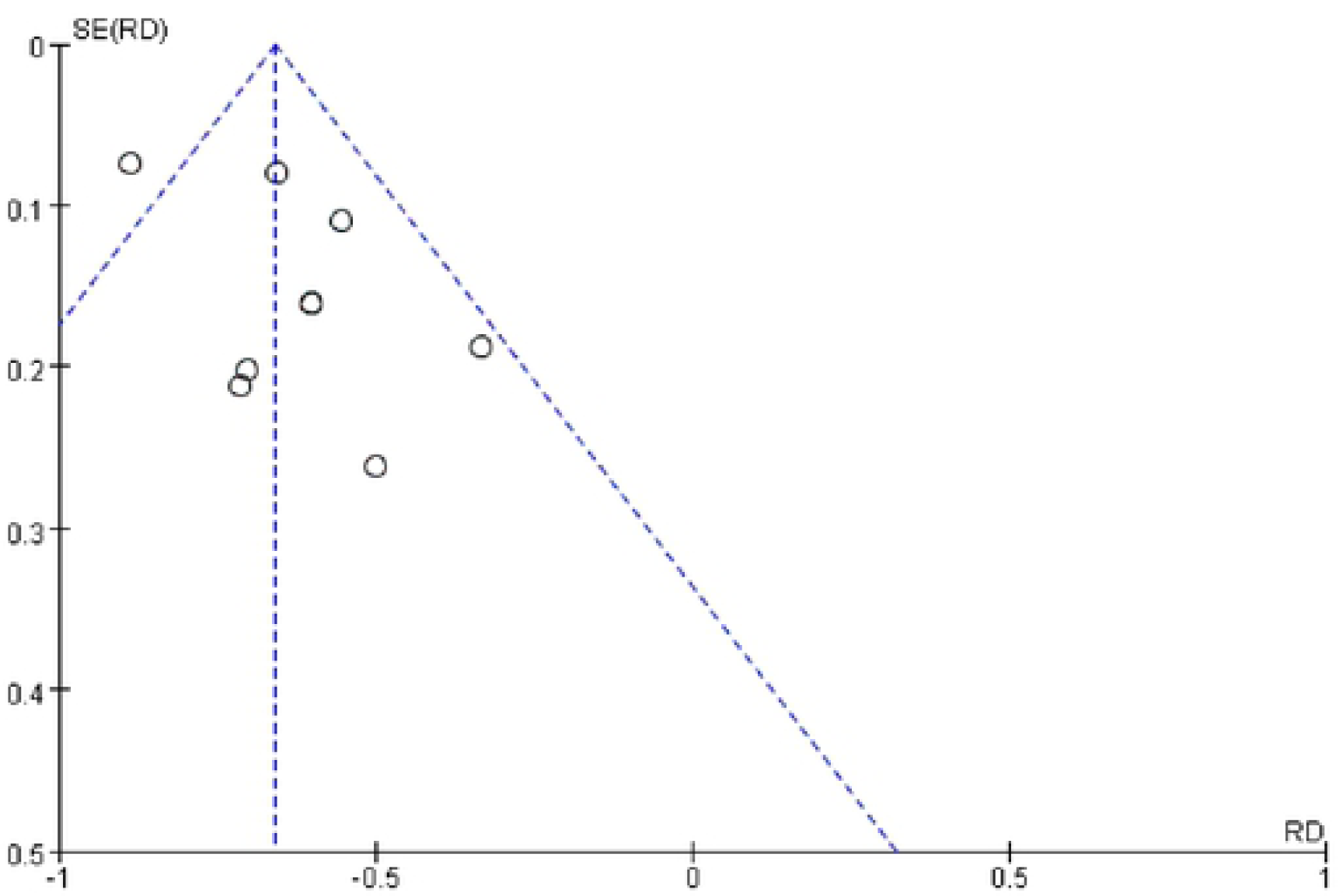
Funnel plots of meta-analysis.

## Discussion

The present meta-analyses attempted to summarize and quantify the immune effect of VP1. The meta-analyses of the protection evidenced an effect with VP1. The benefits of undertaking this meta-analysis clarified the effect of VP1. The analysis showed that VP1 had a protective effect on FMDV [*MH* = -0.66, 95% *CI* = (−0.75, -0.56), *P* <0.00001].

It has been shown that VP1 can induce neutralizing antibodies in experimental and natural hosts^[17]^. VP1 protein and its epitopes have become a research hotspot as a new vaccine and diagnostic product. Some scholars have used the VP1 protein expressed by Hansenula polymorpha to play a certain protective role in the experiment of FMDV in mice and guinea pigs^[18]^. At the same time, some studies have found that VP1 nucleic acid vaccine and virus-like particles expressing VP1 DNA can also play a protective role in FMDV challenge protection experiments^[19]^. The FMDV VP1 protein expressed by Wang and Pan Li in tobacco and tomato plants also showed good immune effects in animal protection experiments^[20, 21]^. Luo et al. used different recombinant plasmids containing VP1 to immunize animals, and they obtained good protection in the FMDV challenge experiment. The specific effect needs further study^[22, 23]^. This previous work established that the nucleic acid or plasmid containing the VP1 gene, as well as various forms of expressed VP1 protein, could provide different levels of protection in immune challenge experiments.

Researchers extensively studied the effects of various FMDV antigens on animal antibodies, but there have not been many studies on challenge protection experiments. The bad results have not been published with *I*^*2*^ = 43%. In the experiment published by Jing et al., it was found that the FMDV VP1 protein expressed by pBAD/TOPO had a certain protective effect in the FMDV challenge experiment of guinea pigs, but the protection rate was relatively low. This may have been because the fusion protein had not been purified. The inclusion body contains many contaminating proteins, and its inclusion body is not completely dissolved. If the expression product inclusion body is further purified or the dose of the vaccine is increased, a better protective effect may be obtained^[24]^. The VP1 expressed in our experiments was not able to effectively induce neutralizing antibodies. This may have been because we used a mouse model, which is insensitive to FMDV and cannot effectively produce antibodies against FMDV.

There is a large amount of research literature available on FMDV vaccines, and meta-analyses can summarize a large number of studies and evaluate experimental results. Researchers could save a lot of reading time by consulting meta-analyses. However, meta-analyses are mainly used in human medicine. There are few applications in veterinary medicine. However, their use in veterinary medicine also has great value. For example, an evaluation of FMD vaccine requires at least $10,000. Analysis of the literature on vaccine evaluation could avoid unnecessary costs. Meta-analysis is a basic statistical method for quantitatively synthesizing previous research data. The advantages are greater statistical power and better resolved inconsistencies between research results. Through meta-analysis, we here established that VP1 has a protective effect on FMDV and the difference is statistically significant. There is less heterogeneity in the meta-analysis of the VP1 experimental group and the control group. The method of endpoint observation might be a source of heterogeneity.

## Conclusion

Through meta-analysis, we here established that VP1 has a protective effect on FMDV and the difference is statistically significant. There is less heterogeneity in the meta-analysis of the VP1 experimental group and the control group. The method of endpoint observation might be a source of heterogeneity. It is necessary to study the epitopes of VP1 to produce new vaccines. VP1 could protect animals from FMDV attacks. It is necessary to study the VP1 protein and its epitopes and use it as a new vaccine and diagnostic product.

## Acknowledgments

We would like to thank LetPub (www.letpub.com) for providing linguistic assistance during the preparation of this manuscript.

